# Narrative Event Segmentation in the Cortical Reservoir

**DOI:** 10.1101/2021.04.23.441090

**Authors:** Peter Ford Dominey

## Abstract

During continuous perception of movies or stories, awake humans display cortical activity patterns that reveal hierarchical segmentation of event structure. Sensory areas like auditory cortex display high frequency segmentation related to the stimulus, while semantic areas like posterior middle cortex display a lower frequency segmentation related to transitions between events (Baldassano et al. 2017). These hierarchical levels of segmentation are associated with different time constants for processing. Chien and Honey (2020) observed that when two groups of participants heard the same sentence in a narrative, preceded by different contexts, neural responses for the groups were initially different and then gradually aligned. The time constant for alignment followed the segmentation hierarchy: sensory cortices aligned most quickly, followed by mid-level regions, while some higher-order cortical regions took more than 10 seconds to align. These hierarchical segmentation phenomena can be considered in the context of processing related to comprehension. Uchida et al. (2021) recently described a model of discourse comprehension where word meanings are modeled by a language model pre-trained on a billion word corpus (Yamada et al 2020). During discourse comprehension, word meanings are continuously integrated in a recurrent cortical network. The model demonstrates novel discourse and inference processing, in part because of two fundamental characteristics: real-world event semantics are represented in the word embeddings, and these are integrated in a reservoir network which has an inherent gradient of functional time constants due to the recurrent connections. Here we demonstrate how this model displays hierarchical narrative event segmentation properties. The reservoir produces activation patterns that are segmented by the HMM of Baldassano et al (2017) in a manner that is comparable to that of humans. Context construction displays a continuum of time constants across reservoir neuron subset, while context forgetting has a fixed time constant across these subsets. Virtual areas formed by subgroups of reservoir neurons with faster time constants segmented with shorter events, while those with longer time constants preferred longer events. This neurocomputational recurrent neural network simulates narrative event processing as revealed by the fMRI event segmentation algorithm of Baldassano et al (2017), and provides a novel explanation of the asymmetry in narrative forgetting and construction observed by Chien and Honey (2020). The model extends the characterization of online integration processes in discourse to more extended narrative, and demonstrates how reservoir computing provides a useful model of cortical processing of narrative structure.

## Introduction

Human existence is embedded in a never ending flow of time (Lashley 1951). A major function of the nervous system is to segment and organize the spatiotemporal flow of perception and action into coherent and relevant structure for effective behavior (Speer et al 2009, Tversky & Zacks 2013, Zacks et al 2007). A particular challenge is to take into account the context of the recent and distant past while addressing the current situation. One of the highest expressions of this is human narrative, which provides a mechanism for encoding, transmitting and inventing reality with its inherent temporal structure (Boyd 2018, Bruner 1991, Ricoeur 1984). Recent advances in neuroscience recording and analysis methods have made possible the study of temporal structure of neurophysiological signals in the human brain during narrative processing, e.g. (Baldassano et al 2017, Chien & Honey 2020, Willems et al 2020).

One of the questions addressed in such research concerns the actual computations that integrate past and present information within the hierarchical networks in the brain (Chien & Honey 2020). Interestingly, networks of neurons that have recurrent connections are particularly well suited for problems that require integration of the past with the present (Dominey 1995b, Elman 1991, Pineda 1987, Servan-Schreiber et al 1991). When we ask how such recurrent networks might be implemented in the brain it is even more interesting that one of the principal characteristics of cortical connectivity is the high density of local recurrent connections (Douglas et al 1995), i.e. that one of the primary characteristics of primate cortex is that it is a recurrent network.

It is thus not surprising that recurrent networks have been used to model cortical processing of sequential and temporal structure, explaining both behavior and neurophysiology (Cazin et al 2019, Dominey 1995a, Enel et al 2016, Laje & Buonomano 2013, Paton & Buonomano 2018). A subset of recurrent network models use fixed (rather than modifiable) random connections which avoids truncating the recurrent dynamics by recurrent learning mechanisms (Pineda 1987). This allows the full expression of recurrent dynamics and high dimension expression of the inputs in the recurrent dynamics. This reservoir computing framework has been used to explain aspects of primate cortical neurophysiology in complex cognitive tasks (Enel et al 2016, Fusi et al 2016, Rigotti et al 2013)

In the domain of narrative and discourse comprehension, Uchida et al (2021) recently used a reservoir-based model to explain how discourse context is integrated on-line so that each new incoming word is processed in the integrated context of the prior discourse. The model addressed the immediacy constraint, whereby all past input is always immediately available for ongoing processing (Hagoort & van Berkum 2007, Just & Carpenter 1980). To address this constraint, the model proposed that making past experience immediately accessible in discourse or narrative comprehension involves a form of temporal-to-spatial integration on two timescales. The first timescale involves the integration of word meaning over extended lifetime, and corresponds to the notion of lexical semantics. This is modeled by the Wikipedia2ec language model (Yamada et al 2020). Wikipedia2vec uses three models to generate word embeddings: a word2vec skip-gram model (Mikolov et al 2013) applied to the 3 billion word 2018 Wikipedia corpus, the link-graph semantics extracted from Wikipedia, and anchor-context information from hyperlinks in Wikipedia (Yamada et al 2020). The second timescale is at the level of the on-line integration of words in the narrative. This is modeled by a recurrent reservoir network that takes as input the sequence of word embeddings corresponding to the successive words in the discourse. The model is illustrated in Figure 1. In (Uchida et al 2021), the reservoir was trained to generate the discourse vector, a vector average of the input words in the discourse. The model was used to simulate human brain responses during discourse comprehension as the N400 ERP, a neurophysiological index of semantic integration difficulty (Kutas & Federmeier 2011). N400 amplitude increases as a function of the semantic distance between a target word and the prior discourse (Metusalem et al 2012, Nieuwland & Van Berkum 2006). This neurocomputational model was the first to simulate immediacy and overrule in discourse-modulated N400.

**Figure 1.**
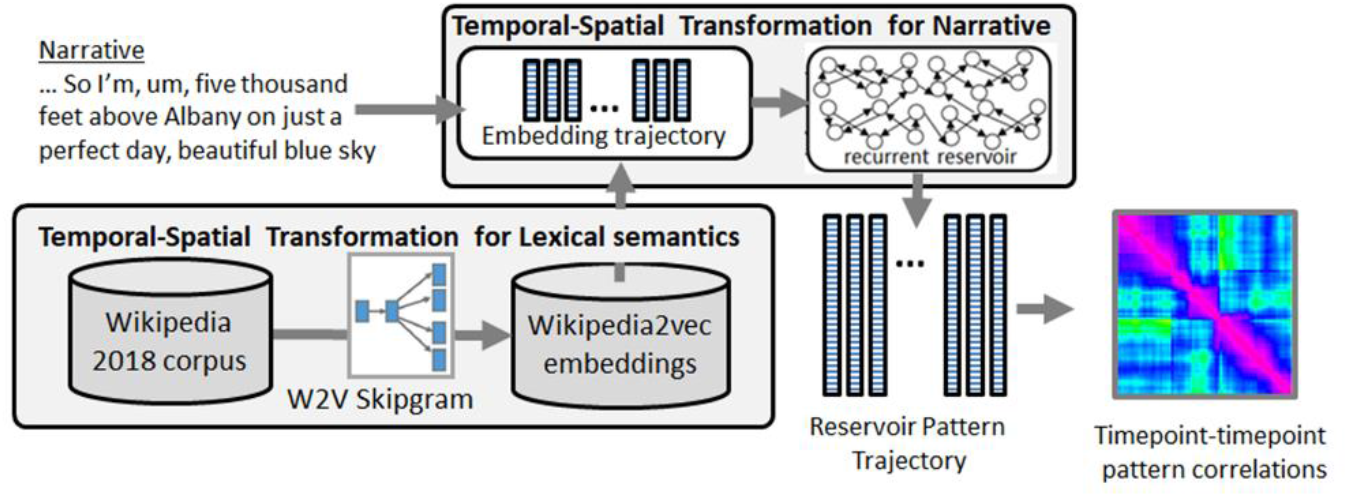
Narrative Integration Reservoir. Word embeddings, generated by Wikipedia2vec, input to reservoir which generates a trajectory of internal states that repreosent the word-by-word processing of the narrative.

The current research explores how this model can account for human neurophysiology of narrative processing in much more extended narratives, increasing from the order of 10^1^ to 10^2^ – 10^3^ words. It is important to acknowledge that this research would not be possible without the open science approach in modern neuroscience and machine learning. Data and algorithms used in the analysis of the human neuroscience are publicly available (Baldassano et al 2017, Nastase et al 2020), which make it possible to use the same analyses on human fMRI and on Narrative Integration Reservoir simulations.

The Narrative Integration Reservoir model embodies two hypotheses: First, that word embeddings from a sufficiently large corpus can serve as a proxy for distributed neural semantics. Second, that temporal-spatial integration of these word vectors in a recurrent reservoir network simulates cortical temporal-spatial integration of narrative. To test these hypotheses the model is confronted with two principal observations related to cortical processing of narrative. The first has to do with the appearance of transitions between coherent epochs of distributed cortical activation corresponding to narrative event boundary segmentation (Baldassano et al 2017). The second has to with a hierarchy of time constants in this processing, and an asymmetry in time constants for constructing vs. forgetting narrative context (Chien & Honey 2020).

Baldassano et al (2017) developed a novel algorithm to detect narrative event boundaries in the fMRI signal of subjects listening to narratives. Their algorithm is a variant of the hidden Markov model (HMM). It identifies events as continuous sequences in the fMRI signal that demonstrate high similarity with an event-specific pattern, and event transitions as discontinuities in these patterns. They used the HMM to segment fMRI activity during narrative processing and made several remarkable observations. In particular they demonstrated that segmentation granularity varies along a hierarchy from short events in sensory regions to long events in high order areas (e.g. TPJ) representing abstract, multimodal situation models.

This allows us to pose the question, are these higher level representations in areas like TPJ the result of longer effective time constants in these higher cortical areas, or is there some processing taking place that is imposing these longer time constants on these higher processing areas? Indeed, in a large distributed recurrent network it is likely that different populations of neurons with different effective time constants will emerge, as demonstrated by Bernacchia et al (2011). We can thus predict that a distribution of neurons with different time constants will be observed in the reservoir and that these time constants will be related to aspects of their narrative segmentation processing.

Interestingly Chien and Honey (2020) demonstrated that the expression of such time constants in narrative processing is dependent on context. In their experiment, one group of subjects listened to an intact narrative (e.g. with a structure ABCD), and another to a narrative that had been scrambled (e.g. with a structure ACBD). The fMRI responses were then compared across these groups in two contexts. The *forgetting* context compared the situation where the two groups started by hearing the same narrative, and then shifted to two different narratives (e.g. AB in group 1 and AC in group 2). Thus the forgetting context was the transition from same (A) to different (B/C). The *construction* context compared the situation where the two groups started hearing different narrative and then began to hear the same narrative (e.g CD in group 1 and BD in group 2). The transition from different (C/B) to same (D) was the construction context. In this clever manipulation of forgetting and constructing context, Chien and Honey (2020) discovered that the time constants of event structure such as observed by Baldassano et al (2017) is reflected in the rate of context construction, whereas there is a uniform rate of context forgetting across these cortical areas. That is, higher cortical areas (e.g. TPJ) construct contexts more slowly, whereas all areas can forget context at a similar rate. In order to account for these results, they developed a hierarchical auto-encoder in time (HAT) model. Each hierarchical level in the model receives prior context from the level below, and higher levels have longer explicit time constants, so their context is less influenced by their input at each time step. This allows the system to account for the hierarchy of timescales in context construction. A surprise signal (that is triggered at event segment boundaries) gates the integration of new input and thus allows for a uniform and rapid reset in the case of context forgetting. These observations on event segmentation, the relation between timescales of processing and segmentation granularity, and the asymmetry in timescales for context forgetting and construction provide a rich framework and set of observations against which we can compare the Narrative Integration Reservoir.

In our experiments, the input to the reservoir of the word vectors for each successive word in the narrative produces a spatiotemporal trajectory of reservoir states. This reservoir activation trajectory simulates the fMRI signal of humans that listen to the same narrative, and we can thus apply the segmentation HMM of Baldassano et al (2017) to these reservoir state trajectories.

To evaluate the Narrative Integration Reservoir, we first expose it to a set of short texts extracted from the New York Times, and Wikipedia to test whether the reservoir will generate activity patterns that can be segmented by the HMM. Next we compare reservoir and human brain neural activity trajectories generated by exposure to the same narrative. We then undertake two more significant tests of the model. First, we examine whether the model can provide insight into the asymmetry of construction vs forgetting of narrative context a observed by Chien and Honey (2020). We then test the hypothesis that different effective time constants for reservoir neurons will be associated with different granularity of event segmentation as observed in fMRI data by Baldassano et al (2017).

## Results

### Segmentation at topic boundaries

We first tested the hypothesis that the Narrative Integration Reservoir when driven by narrative input should exhibit event-structure activity with HMM segmentation correlated with known narrative boundaries. We created two test narratives with clearly identifiable ground truth boundaries. For the first, we choose four short text segments from different articles in the New York Times and concatenated these together to form a single narrative text that had well defined topic boundaries. We performed the same operation with four short text segments from 4 Wikipedia articles to generate a second narrative with known ground truth boundaries. For the Wikipedia test narrative, the word length of the four constituent texts were quite different, with the first text longer than the sum of the other three. This yielded our two test narratives which were then used as input to the Narrative Integration Reservoir in order to generate the trajectory of reservoir activation. We tested the resulting event-structured activity by applying the HMM segmentation model of Baldassano et al (2017) to the trajectory of reservoir states generated by feeding the embeddings for the words in these texts into the reservoir. The HMM takes as input the neural trajectory to be segmented, and the number of segments, K, to be identified, and produces a specification of the most probable event boundaries. The HMM model is available in the python Braniak library described in the methods section. The results are illustrated in Figure 2.

**Figure 2.**
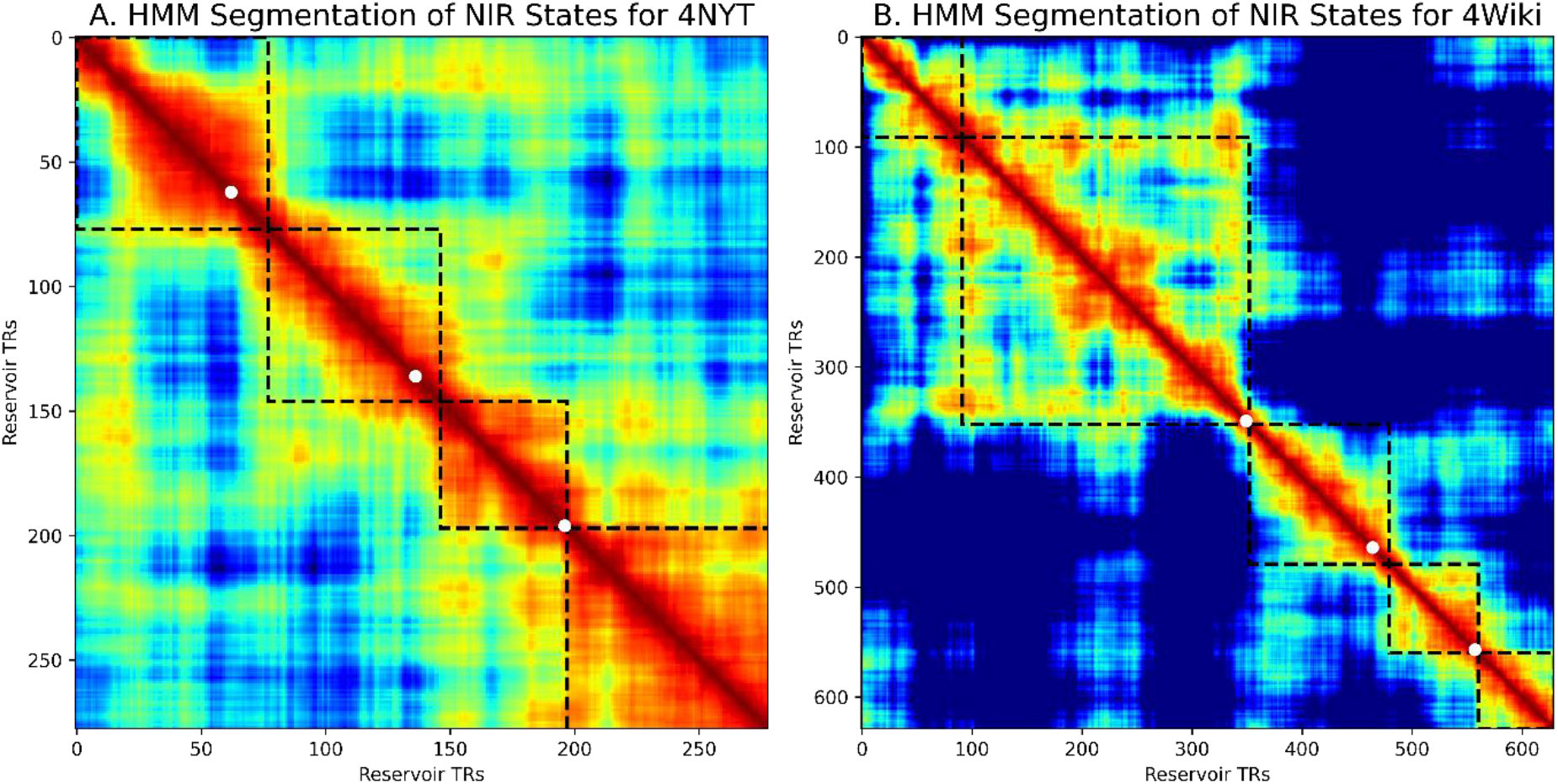
Time-point time-point corrlations of reservoir activity during processing of input narratives. Left 282 word text made of four paragraphs from different articles in the New York times. Black dotted lines indicate segmentation of the HMM. White points indicate ground truth section boundaries in the text. Right Longer input txt with greater variability in section lengths. Same notation as right. HMM segmentation reasonably compares to ground truth.

The left panel illustrates the timepoint-timepoint correlations for reservoir activity during processing of the 278 word four-segment New York Times narrative. Coherent zones correspond to the four distinct text segments. The dotted lines indicate event boundaries identified automatically by the HMM with pre-specified K = 4, and white dots indicate the actual ground truth boundaries. The narrative boundaries marked by the white dots are closely followed by the event boundaries discovered by the HMM. The right panel similarly illustrates automatic and ground truth boundaries for 628 word Wikipedia test narrative. In this case there is an additional boundary detected within the first text, with pre-specified K = 5.

To evaluate the segmentation, we first compared the segment boundaries discovered by the HMM with the ground truth values (i.e. the segment boundaries between the four paragraph of each text). The ground truth and HMM segment boundaries are highly correlated, with the Pearson correlation r = 0.99, p < 0.0001. To evaluate the specificity of the segmentation for each of the two texts, we segmented both under the same conditions, with pre-specified K=5. We then normalized the resulting event boundaries into a common range, and made a pairwise comparison of the segment boundaries for the two texts. The event segmentation discovered by the HMM for the New York Times text is clearly different from that for the Wikipedia text (t(5)=−4.9, p < 0.05). This allows us to consider that the Narrative Integration Reservoir can display some form of event segmentation that is specific to the given text to which it is exposed. Given this first indication of segmentation capability, we can proceed with more detailed analyses.

### Comparison of Narrative Integration Reservoir and Human fMRI Segmentation on same input

Baldassano et al (2017) demonstrated cross-modal segmentation, where application of their HMM to fMRI from separate groups that either watched a movie or listened to someone recalling the movie produced similar event segmentation. Analogously, we set out to compare HMM segmentation in two conditions: The first is recurrent reservoir activity when the Narrative Integration Reservoir is exposed to the text transcript of an auditory story. The second is the fMRI signal from humans who listened to an auditory version to the same story.

Thanks to their open data policy (Baldassano et al 2017, Chien & Honey 2020, Nastase et al 2020), we have access to fMRI from human subjects that listened to an audio narrative about a skydiving lesson called “It’s Not the Fall”, along with a transcript of this narrative. We can thus use the transcript to generate a sequence of word embeddings from Wikipedia2vec which is the input to the Narrative Integration Reservoir. The resulting trajectory of activation patterns in the reservoir can be segmented using the HMM. In parallel the HMM is used to segment human fMRI from subjects who listed to the same story. This allows a parallel comparison of the HMM segmentation of human brain activity and of Narrative Integration Reservoir activity, as illustrated in Figure 3.

**Figure 3.**
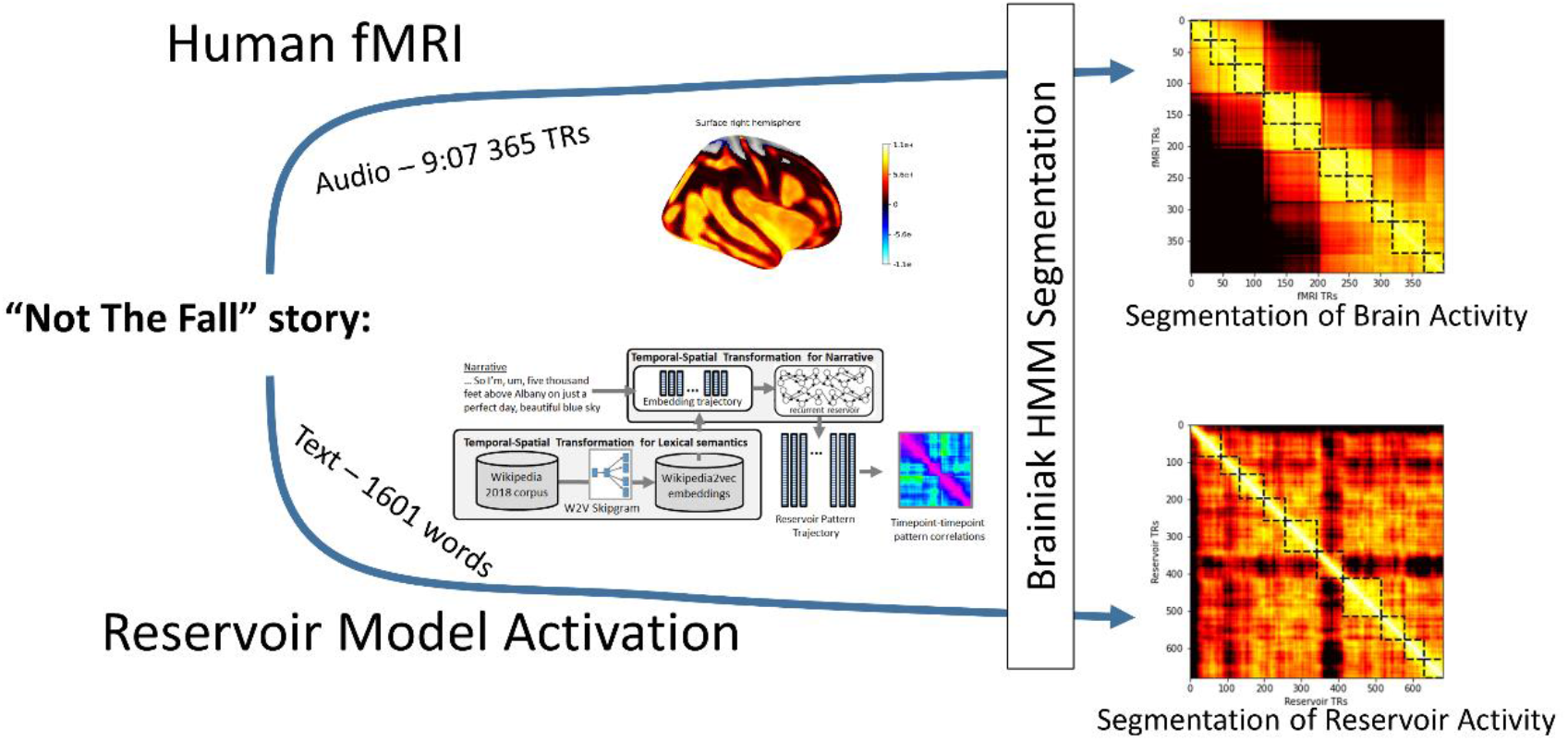
Pipeline for comparting HMM segmentation of human fMRI and reservoir recurrent states during processing of the same narrative. Humans listen to the “Not The Fall” story in fMRI scanner. Narrative Integration Reservoir model exposed to transcript of the “Not The Fall” story. Resulting trajectories of human fMRI and model reservoir states are processed by the Baldassano HMM.

Data from 10 subjects were compared with state trajectories from 10 reservoirs. The subjects listened to the audio narrative “It’s not the fall that gets you” (Nastase et al 2020) which was used in Chien and Honey (2020). This is a story told by a comedian of his skydiving experience, with a young guide who teaches him to land (because it’s not the fall that gets you, it’s the landing!), and who also plays a trick on him saying there is something wrong with the chute as he jumps from the plane. The audio had a duration of 9:47 and this corresponds to a total recording of 365 TRs. The fMRI data contains 400 TRs corresponding to an initial 22 seconds of music, 3 seconds of silence, the story, and then 10 seconds of silence after the story.

For each subject, the HMM was run on down-sampled whole brain neural activity with pre-specified K = 10. At each TR, the whole brain image of 110592 voxels was down sampled to 10000 voxels. Note that we do not require detailed analysis by region of interest, but rather a general global effect. The summary results of this segmentation are presented in Figure 4A. There we see the segmentation into 10 events in the average over all subjects. Likewise for the Narrative Integration Reservoir, 10 instances were created using different seed values for initialization of the reservoir connections, and each was exposed to the word by word narrative. For each word, the corresponding Wikipedia2vec 100 dimensional embedding was retrieved and input into the reservoir. The HMM was then run on each of these reservoir activation trajectories. The summary results of this segmentation are presented in Figure 4B.

**Figure 4.**
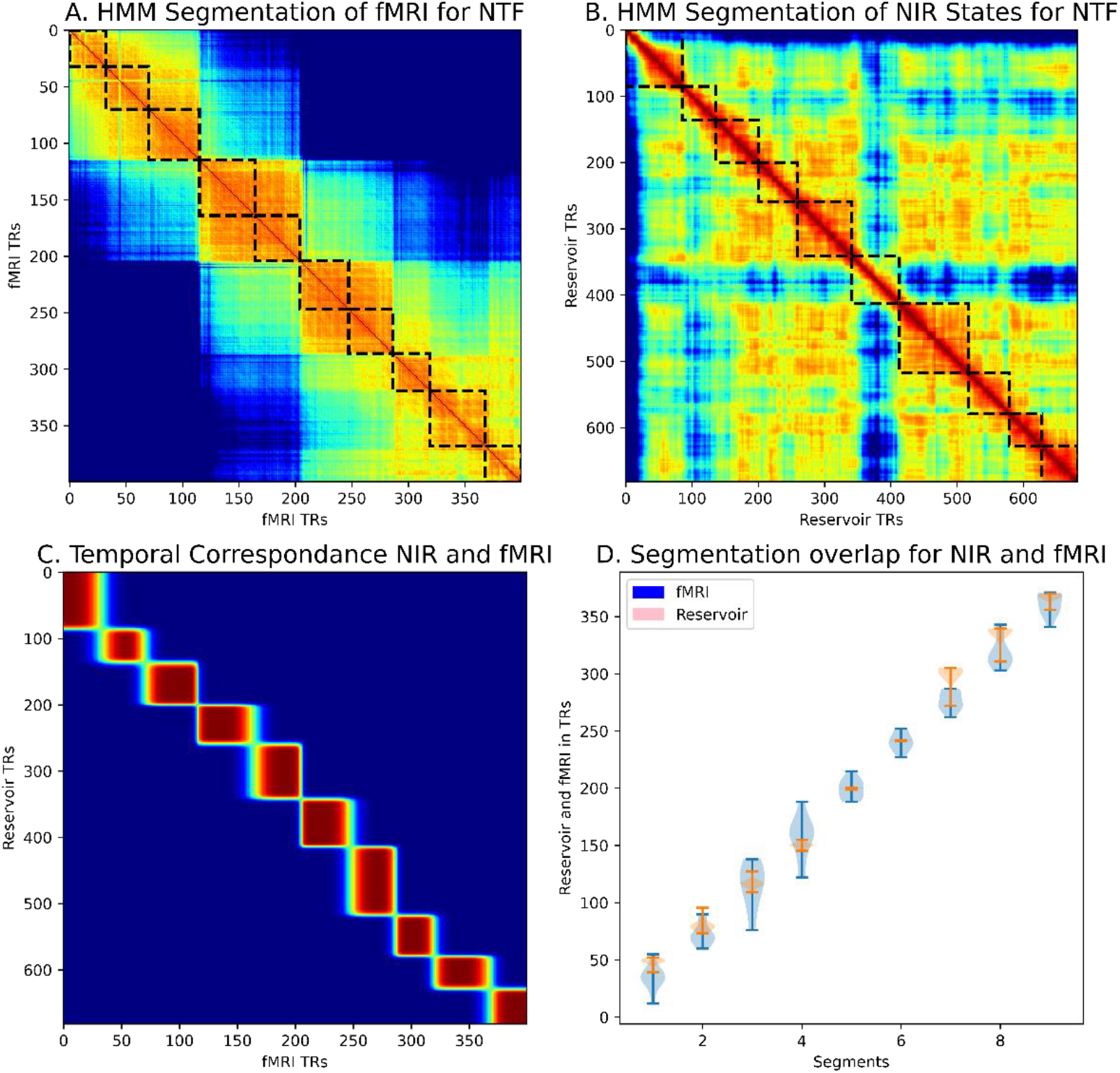
Human and Model HMM segmentation. A: Segmentation of human fMRI. B: Segmentation of Narrative Integration Reservoir model internal states. C: Temporal correspondence of fMRI and model segmentation. D: Correspondence of segmentation boundaries for fMRI from 10 subjects (blue) and reservoir trajectories from 10 model instances (pink).

In order to visualize the temporal correspondence between the reservoir and the fMRI, we can calculate the probability that reservoir state and an fMRI TR are in the same event (regardless of which event it is). This is

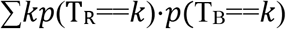

which we can compute by a simple matrix multiplication on the two segmentation matrices^1^. The result is illustrated in Figure 4C which illustrates the temporal correspondence between the segmentation of human fMRI and Narrative Integration Reservoir state trajectories. We further undertook two methods to determine the correspondence between events detected in the fMRI signal of humans listening to the narrative, and in the Narrative Integration Reservoir state trajectory signal of reservoirs exposed to the sequence of word embeddings generated from the transcript of the narrative. The first method is to compare the timing of the segmentation boundaries in a common timeframe. We used the fMRI TR time frame. To convert reservoir time steps to fMRI TRs we divided by the total reservoir time steps and multiplied by the total number of fMRI TRs. Then for each human subject, we collected the HMM segment boundaries for the discovered segments, and the same for each of the tested reservoirs. The distributions of segmentation boundaries are displayed in Figure 4D. There we observe that for all of the boundaries, the human and model distributions overlap. There is a significant correlation between the segmentation boundaries for the human fMRI and Narrative Integration Model, Pearson correlation r = 0.99, p < 0.0001.

The second method to determine the correspondence between human and model event boundaries was to actually go back into the two narrative sources (the human auditory narrative and the model transcript) and compare the words in the auditory and text narratives at the boundaries. For the human data, the 9 segment boundaries in TRs were converted to seconds (1.5sec/TR), and then the appropriate context was identified at the corresponding time in seconds in the audio file that subjects listened to^2^. Likewise, for the reservoir data, the 9 segment boundaries in reservoir time steps correspond to the numerical input order of the word embeddings, and so the corresponding words were identified. Interestingly, 7/9 of the identified segments in the human and reservoir data correspond to the same salient events in the story. These include (1) driving upstate to the skydiving school, (2) signing the contract, (3) the “drop and roll” landing procedure, (4) progressively raising the practice landing/diving platform, (5) fear of chute not opening and the “last couple of minutes”, (6) after landing, anger and “what the heck was that!”, (7) heart starts slowing, realizing he survived. Two of the unpaired events corresponded to audience laughing in the audio that was not present in the transcript. While this second method is not directly amenable to a detailed statistical analysis, it does allow us to appreciate that meaningful events detected in the fMRI signal were also detected in the Narrative Integration Reservoir.

The important point is that the Narrative Integration Reservoir demonstrates structured representations of narrative in terms of coherent trajectories of neural activity that are discontinuous at event boundaries, as revealed by the HMM segmentation. This allows us to proceed with investigation of temporal aspects of this processing.

### Different Timing of Constructing and Forgetting Temporal Context

In order to investigate the time course of context processing, we exposed the reservoir to an experimental manipulation based on that used by Chien and Honey (2020). We recall that they considered two types of contextual processing: constructing and forgetting. In the constructing context, separate groups of subjects heard different narratives and then at a given point began to hear the same narrative. At this point, they began to construct a shared context. Conversely, in the forgetting context, two separate groups of subjects initially heard the same narrative, and then at a given point began to hear two different narratives. At this point, they began to forget their common context.

We thus exposed paired instances of the Narrative Integration Reservoir (i.e. two identical instances of the same reservoir) to two respective conditions. The first instance was exposed to an intact version of the Not the Fall transcript in four components ABCD. The second instance was exposed to a shuffled version of the transcript ACBD.

For the two model instances, the initial component A is the same for both. The second and third components BC and CB, respectively, are different, and the final component D is the same. We can thus examine the transition Same to Different for forgetting, and the transition Different to Same for constructing. We expose two identical copies of the same reservoir separately to the intact and shuffled conditions, and then directly compare the two resulting reservoir state trajectories by subtracting the shuffled from the intact trajectory. This is illustrated in Figure 5. There we see that in the common initial Same section, there is no difference between the two trajectories. At the Same to Different transition we see an abrupt difference. This corresponds to forgetting the common context. Then in the transition from Different to Same, we see a more gradual convergence of the difference signal to zero.

**Figure 5.**
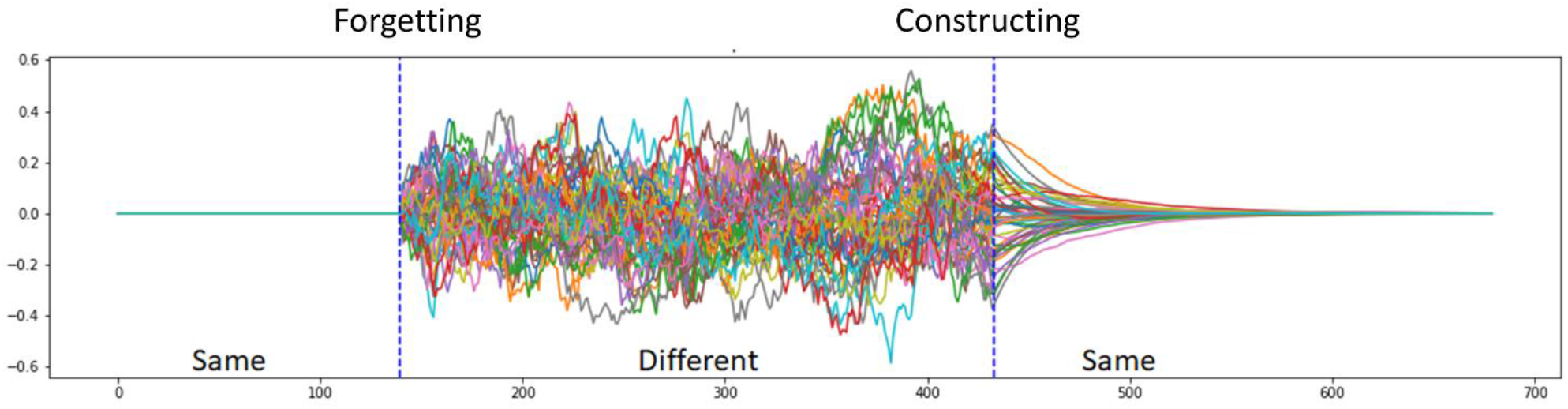
Reservoir activity differences for two Narrative Integration Reservoirs exposed to the Same-Different and Different-Same transitions. In the initial Same period, both perceived the same initial 140 words of the “Not The Fall” story. In the following Different period, model 1 continued with the story, and model 2 received a shuffled version. In the final Same period, both were again eposed to the same final portion of the story. Forgetting, at the Same-Different transition, takes place relatively abruptly. Constructing, at the Different-Same transition appears to take place much more progressively.

To analyse construction and forgetting across the two groups, Chien and Honey (2020) measured the inter-subject pattern correlation (ISPC) by correlating the spatial pattern of activation at each time point across the two groups. We performed the same correlation analysis across the two reservoirs. In Figure 6 we display the forgetting and constructing context signals, along with the pairwise correlation diagrams, formed by the correlation of states at time t of the intact and shuffled reservoir trajectories. Gradual alignment or constructing is illustrated in Figure 6A which shows the difference between reservoir trajectories for intact vs. shuffled inputs at the transition from Different to Same. In Figure 6B the cross-correlation between the intact and shuffled reservoir state trajectories is illustrated, with a blue circle marking the point illustrated by the dotted line in 6A, where the shuffled and intact narratives re-converge to the common Same ending. Interestingly, we see that there is a gradual smooth reduction of the differences in 6A, and progressive re-construction of correlation along the diagonal in 6B. In contract, in 6C and D we focus on the transition from Same to Different. There we see the difference signal and the cross-correlation map for the divergence/forgetting context in the Same to Different transition. Presented in the same timescale as A and B, we see in C and D a much more abrupt signal change, both in the difference signal (6C) and in the cross-correlation map (6D). There, where the white circle marks the shift from common to different narrative input, we see an abrupt end to the correlation indicated by the initially high value along the diagonal which abruptly disappears. Interestingly this corresponds to the observation of Chien and Honey (2020), who found a more extended time constant for constructing vs. forgetting a context. We now examine these effects in more detail.

**Figure 6.**
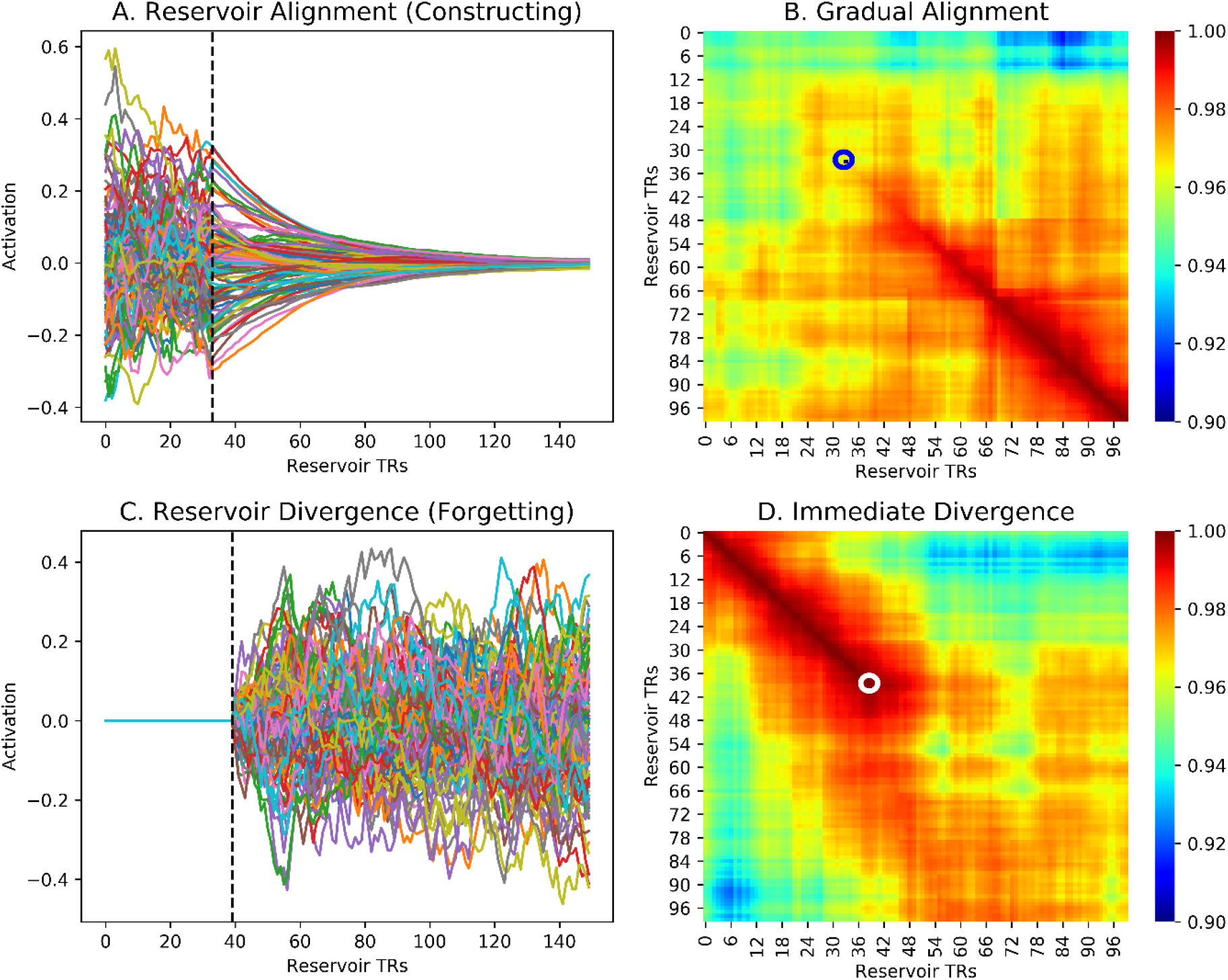
Dynamics of Constructing and Forgetting. A. Zoom on reservoir activity difference during contructing (transition from Different to Same). Dotted line mars the transition. Note the gradual decay. B. Time-point time-point correlations between the two (intact and shuffled input) reservoirs. Blue circle marks the beginning of the Same input. Note the slow and progressive buildup of coherence revealed along the diagonal. C. Zoom on reservoir activity difference during forgetting (transition from Same to Different). Dotted line mars the transition. Note the abrupt increase. D. Same as B, but white circle marks the Same to Different transition. Not the abrupt loss of coherence along the diagonal.

### Distribution of Time Constants for Context Alignment/Construction

In Chien and Honey (2020) this extended time constant for constructing was observed particularly for cortical areas higher in the semantic processing hierarchy such as the temporal-parietal junction TPJ, vs. primary auditory cortex. Indeed, they demonstrated that there is a distributed continuum of time constants across cortex.

Considering such properties in the Narrative Integration Reservoir, we recall that the reservoir is made up of leaky integrator neurons with a leak rate, and thus the reservoir should have some characteristic inherent temporal dynamics and time constants. More importantly, within the reservoir, combinations of recurrent connections can construct subnetworks that have different effective time constants. This predicts that we should be able to identify a distribution of time constants within the reservoir, similar to the observations of Bernacchia et al (2011). To test this prediction, we again exposed paired reservoir instances to the intact and shuffled conditions, and analyzed the difference between the two reservoir state trajectories. For each neuron in the reservoir pair, we took the absolute value of its activation difference (intact – shuffled) at the onset point of the convergence/construction period (indicated by the dotted vertical line in Figure 6A) and counted the number of time steps until that value fell to ½ the initial value. This was used as the effective time constant of the construction rate for each neuron. The sorted values of these time constants is presented in Figure 7 where it compares well with the same type of figure illustrating the distribution of cortical construction time constants from Chien and Honey (2020) Figure S2.

**Figure 7.**
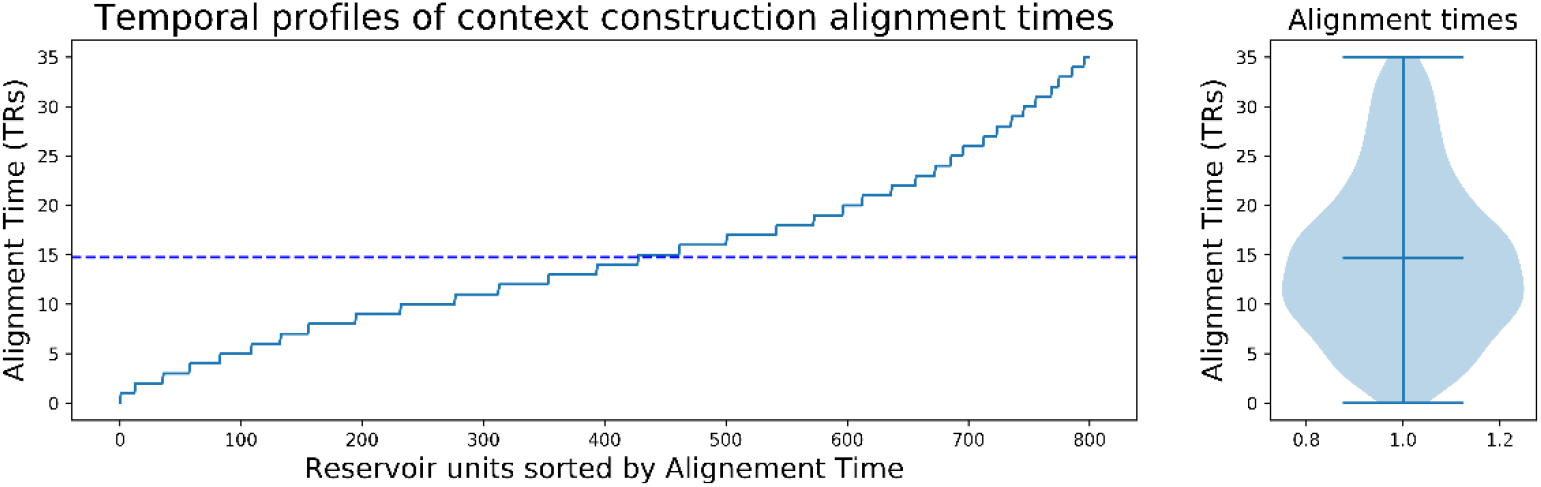
Reservoir units sorted by alignment time. This distribution is remarkably similar to that observed by Chien and Honey (2020) (see their Figure S4B).

We can further visualize the behavior of this distribution of time constants by binning neurons into time constant groups, forming different virtual cortical areas. Figure 8 illustrates five thus created virtual areas of 200 neurons each, from the fastest (left) to slowest (right) time constants. Note the slopes of the neuronal response which become successively more shallow. Below the traces of neural responses, the corresponding cross-correlation between the intact and shuffled reservoir state trajectories for each of these sub-groups of neurons is illustrated. We observe that for the faster neurons, the alignment is faster as revealed by the coherent structure along the diagonal. Figure 9 illustrates these response functions for these five areas in the constructing and forgetting contexts. There we observe that while the time constant for forgetting (transition from Same to Different) is fixed across these five cortical areas, there is a clear distinction in the time course of constructing (transition from Different to Same), corresponding to the observations of Chien and Honey (2020). Now we consider the functional consequences of this diversity of time constants.

**Figure 8.**
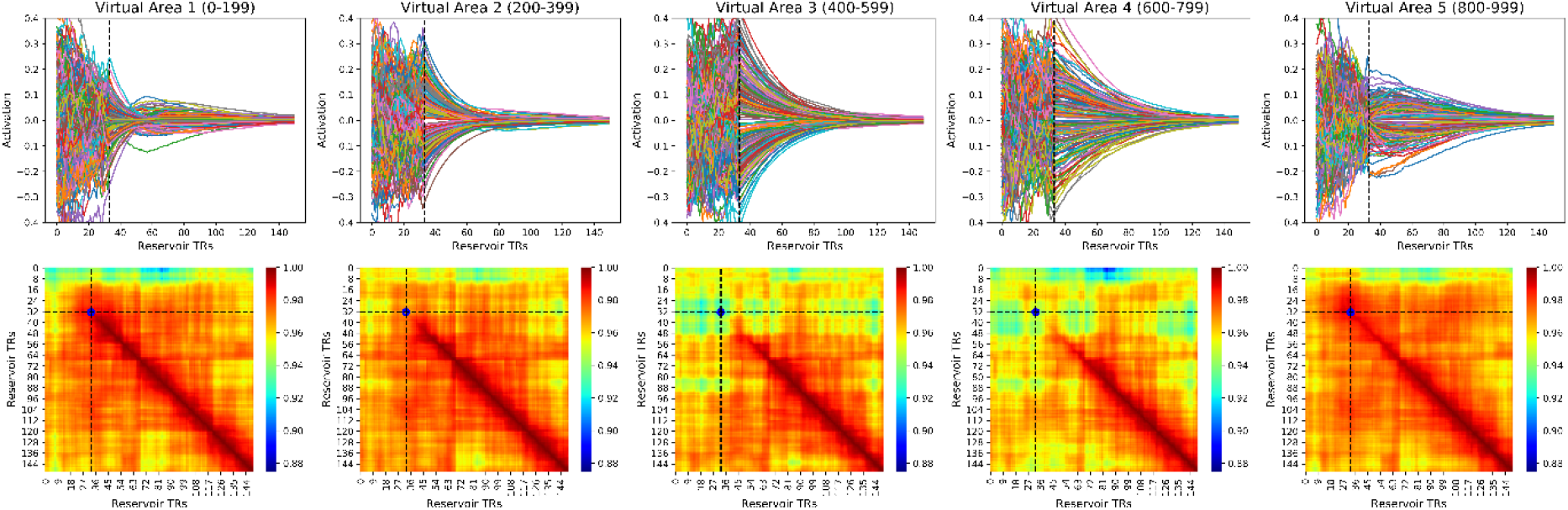
Above - Reservoir unit activity during alignment or constructing after the Different to Same transition, grouped by alignment time. Alignment time increases from left to right in the 5 panels. Note the steep slope of the activity in the leftmost panel the progressively flattens in the successive panels on the right. Likewise note the “rebound” in the leftmost panel, where neurons quickly reduce their activity but then continue and cross the x-axis before coming to zero. In the rightmost panel, corresponding to the slowest neurons, activity actually increases at the transition before slowly coming back to zero. This indicates the complex and non-linear characteristics of the reservoir. Below – Effects of alignment time on context construction. Same format a Figure 68. Note how the onset of coherence along the diagonal varies with the alignment time in the five sets of neurons grouped by alignment time.

**Figure 9.**
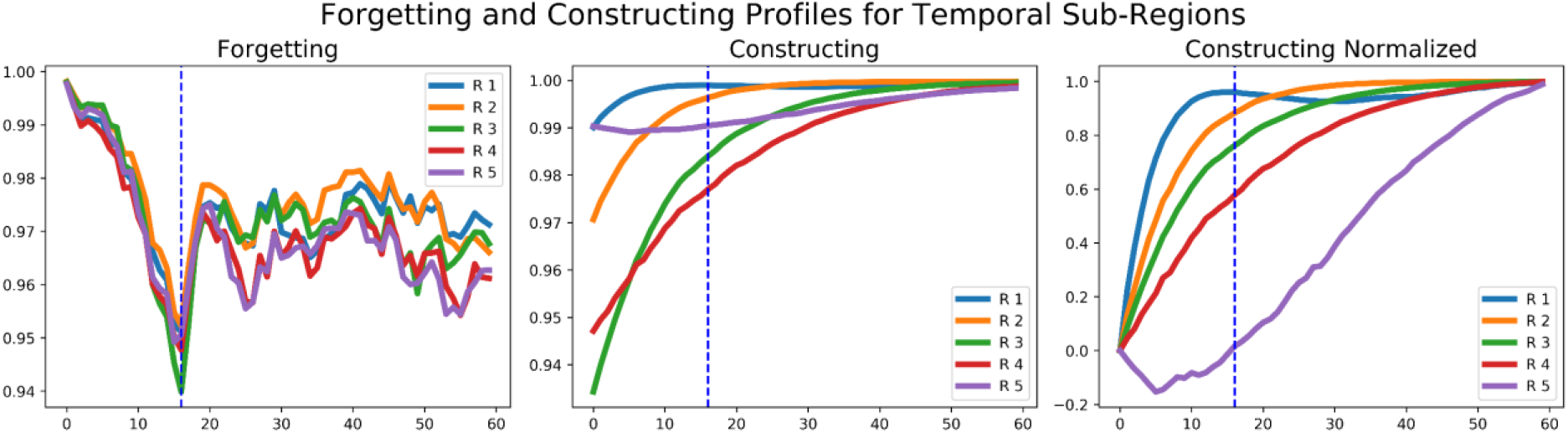
Temporal profiles of Forgetting and Constructing, as revealed by inter-subject pattern correlations, for the 5 temporal groups illustrated in Figure 8. Forgetting. All 5 regions (R1-R5) display an overlapping descent to the minimum coherence in parallel. Constructing. In contrals the 5 regions display a diversity of time-courses in their reconstruction of coherence. Dotted blue line indicate the same time point in both panels. On the right, we display the normalized inter-subject pattern correlations, which make more apparent the different temporal profiles for constructing a coherent representation in the transition from Different to Same for the 5 temporal regions in the reservoir.

### Segmentation granularity corresponds to construction time constant

Baldassano et al (2017) showed that segmentation granularity varies across cortex, with shorter events in sensory areas and increasingly longer events in higher cortical areas. Chien and Honey (2020) further revealed that the time constant for context constructing similarly increases along this hierarchy. We can now test the hypothesis that there is a relation between the time constant for construction, and the granularity of segmentation in the Narrative Integration Model by the HMM. In other words, longer time constants for construction will correspond to a preference for fewer, longer events in segmentation by the HMM, whereas shorter time constants will have a preference for more numerous and shorter events in the segmentation by the HMM.

In order to test this hypothesis, we measure transfer performance of the HMM in coherent conditions, where it is trained and tested on cortical regions that have similar and coherent time constants, and then we compare this with an incoherent condition, where training and testing take place on cortical regions with different and incoherent time constants. In more detail, we run the forgetting-construction experiment as described above, and then generated 10 virtual cortical areas of 100 neurons each, sorted along the construction time constant gradient, so each successive area has an increasingly higher construction time constant as illustrated in Figure 8. We then use the HMM with K = 24 for the 5 fast areas, and K = 8 for the 5 slow areas. The so-called fast HMM (K=24) was trained on a Fast area (neurons 100-199) and tested on the remain 4 fast areas. The so-called slow HMM (K=8) was trained on a Slow area (neurons 800-899) and tested on the other four slow areas. The Under these conditions we expected the trained HMMs to generalize better than in the incoherent condition. The incoherent condition was almost the same as the coherent condition, except that the K values (i.e. number of segments to be found) were reversed, so that fast areas were trained and tested with a slow HMM (K=8), and slow areas with a fast HMM (K=24). We tested this procedure on 100 reservoirs. The result of the experiment revealed that there was a highly significant difference in the transfer (measured as the log likelihood of the model fit), with an increased transfer quality for the coherent vs. incoherent conditions (t=4.9, p = 4.9e-06). Figure 10 illustrates the HMM segmentation results for the fast and slow virtual areas. The left panel A shows the time-point time-point correlation map for the average of 100 fast areas (neurons 100-199) with small convergence time constants, and the event boundaries found by the HMM with K = 24. The right panel B illustrates the same for the average of 100 slow areas (neurons 800-899) with large convergence time constants, and the event boundaries found by the HMM with k = 8. We can observe the finer grained correlation structure along the diagonal in panel A, and more coarse grained, larger event structure in panel B.

**Figure 10.**
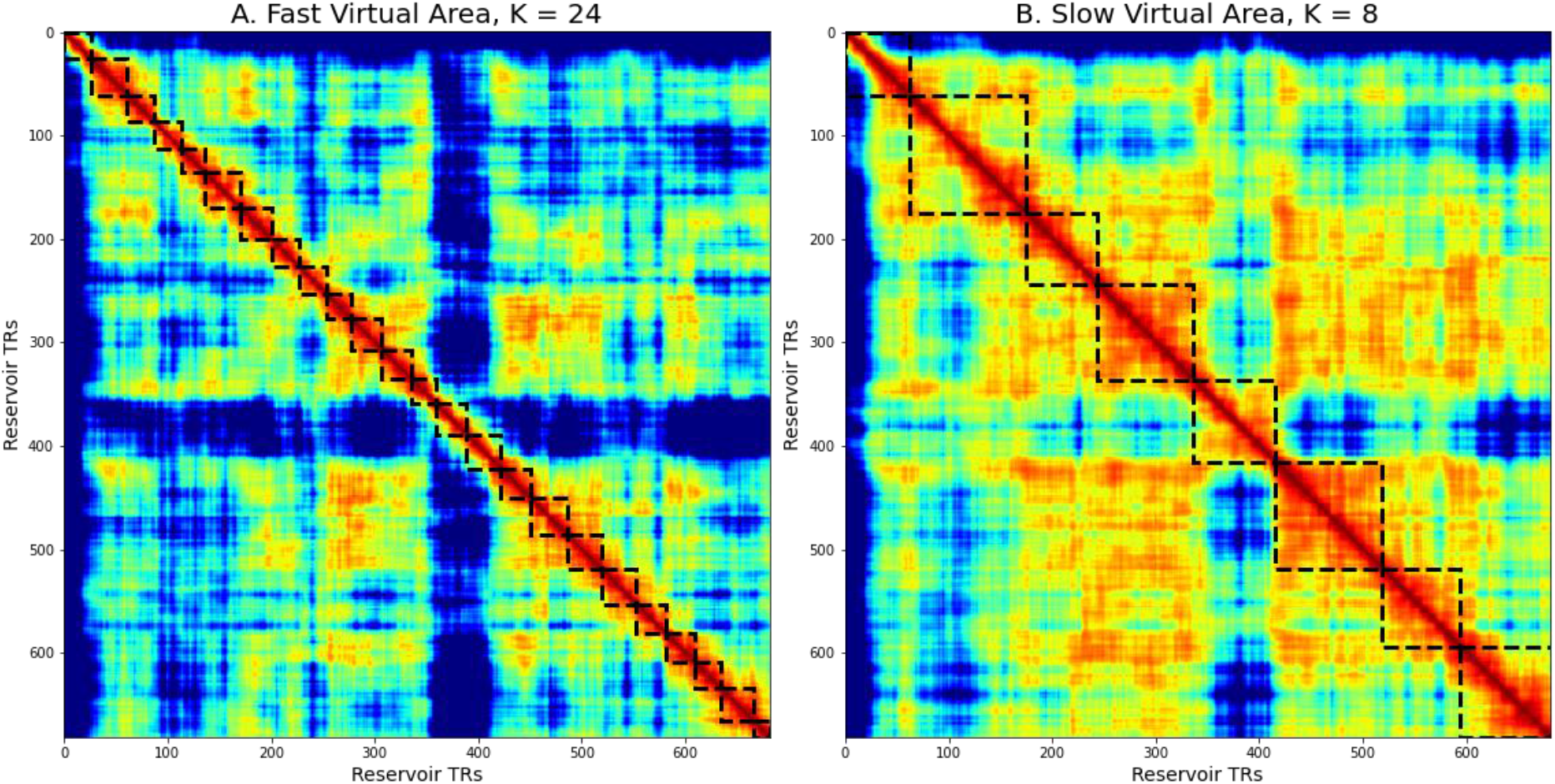
HMM segmentation in two populations of reservoir neurons with faster vs. slower alignment times. A – reservoir subpopulation with shorter alignment times. HMM segmentation into K = 24 event segments. B – reservoir subpopulation with greater alignment times. HMM segmentation into K = 8 regions. Note the larger sections of continuous coherence in the left vs. right panel.

## Discussion

Narrative is a uniquely human form of behavior that provides a coherent structure for linking events over time (Bruner 1991, Polkinghorne 1988). A given instance of a narrative itself is a temporal sequence that must be processed by appropriate mechanisms in the nervous system (Willems et al 2020). Previous research has identified constraints on this processing related to immediacy (Hagoort & van Berkum 2007). Information that is provided early in the narrative must remain immediately accessible at any time. This is technically an interesting problem, as in many architectures, the time to access previous memories may depend on the size of the memory, or the position of the data in question in the input sequence. We recently reasoned that performing a temporal-to-spatial transformation can solve this problem, provided that there is a parallel readout capability that provides immediate access to the distributed spatial representation (Uchida et al 2021).

The current research further tests this model in the context of temporal processing constraints identified in more extended narrative processing. Chien and Honey (2020) characterized the temporal processing of the construction and forgetting of narrative context. Interestingly they observed that whereas across cortical areas, forgetting occurs with fixed timing, construction takes place with a time constant that significantly varies across cortical areas. In order to account for the hierarchy of time constants across areas they developed the HAT model with explicit hierarchically organized modules, where each successive module in the hierarchy is explicitly provided a longer time constant. They introduced a gating mechanism in their hierarchical recurrent model in order to explicitly address the more abrupt response to change in context in the forgetting context. This model is a significant achievement as it provides an explicit incarnation of the processing constraints in the form of a hierarchy of modules, the pre-specified time constants and the explicit gating mechanism. As stated, the problem remains, “what are the essential computational elements required to account for these data?” (p. 681-682). The third constraint comes from Baldassano et al (2017) who demonstrate that higher elements in the cortical hierarchy prefer segmentation with fewer and larger segments, while lower sensory areas prefer more and shorter segments. Part of the motivation for the current research is to attempt to respond to the question “what are the essential computational elements required to account for these data?” with respect to these three constraints.

Indeed, when we simulated the experimental conditions where subjects heard the same and then different narratives (the forgetting context) and respectively different and then same narratives (the constructing context), the between reservoir correlations displayed an abrupt response in the forgetting context, and a distribution of gradual responses in the constructing context. Interestingly, this behavior is an inherent property of the reservoir: there are no specific mechanisms necessary. The hierarchy of decay time constants is a natural property of the reservoir, as observed by Bernacchia et al (2011) (and see Methods below). The transition from different to same requires the dissipation of the past differences in face of common inputs which takes time similar to the response to an input described in the methods section below. In contrast, in the transition from same to different, the rapid onset response in the forgetting context is due to immediate responses to input in the reservoir. Even if the change in each neuron takes place progressively, the difference between the two input driven responses is apparent immediately. This behavior of the reservoir addresses the first two constraints from Chien and Honey (2020) – the construction, forgetting asymmetry, and the hierarchy of time constants across cortical areas for forgetting.

By partitioning the reservoir neurons into subgroups based on their constructing time constants, we observed that areas with longer time constants preferred segmentation with fewer, longer events, while on the opposite, areas with shorter time constants preferred segmentation with more numerous and shorter events, thus addressing the third constraint, from Baldassano et al (2017). While there are likely multiple mechanism that contribute to the diversity of inherent time constants across cortical areas (Murray et al 2014), the internal dynamics of recurrent loops within the reservoir account for a rich diversity of effective time constants as we observe here. We can propose that this segregation is a natural product of neuroanatomy, and that narrative structure has found its way into this structure. Narrative structure reflects the neuroanatomy.

It is worth noting, in the context of the relation between narrative structure and recurrent network functional neuroanatomy, that the observed reservoir behavior comes directly from the input-driven dynamics of the reservoir. There is no learning, and we do not look at the readout, but at the recurrent elements of the reservoir itself.

A limitation of the current modeling effort is that it does not explicitly address meaning. Indeed, in our previous work (Uchida et al 2021), we predicted N400 responses using the trained readout of the reservoir to generate the cumulative average of the input word sequence, thus forming a discourse vector. This could then be compared with the word embedding for a target word in order to predict the N400 as 1-similarity. Related models have used recurrent networks to integrate meaning over multiple word utterances in order to predict the N400 (Brouwer et al 2017, Rabovsky et al 2018), further supporting the role of recurrent connections in accumulating information over multiple words. We have also started to address how structured meaning in the narrative is extracted to build up a structured situation model (Mealier et al 2017, Pointeau et al 2021), so that the system can learn to answer questions about perceived events and narrative.

In the current modeling effort we do not train the reservoir, as there is no explicit task, except to process the sequence of word embeddings corresponding to the narrative. The learning has already occurred, in creation of the Wikipeida2vec embeddings (Yamada et al 2020), which represents sufficient knowledge to allow our system to perform simple human tasks requiring inference on events as we demonstrated in (Uchida et al 2021). Thus, in the current research it is the internal dynamics of the reservoir itself, driven by the Wikipedia2vec embeddings for the successive words in the narratives - that is the object of our analysis – analogous to the internal cortical dynamics analysed in the human fMRI studies (Baldassano et al 2017, Chien & Honey 2020). The degree of resemblance that we found between cortical and reservoir dynamics in narrative processing provides further support for the idea that aspects of cortical function can be considered in the context of reservoir computing (Barak 2017, Enel et al 2016, Fusi et al 2016, Rigotti et al 2013). Future research might examine the presence of high dimensional coding and mixed selectivity, characteristic of reservoir computing, in cortical processing of narrative.

## Material and Methods

### Narrative Integration Reservoir

The end-to-end functioning of the Narrative Integration Reservoir is illustrated in Figure 1. The model consists of two components. The first generates word embedding vectors, and the second generates the spatiotemporal trajectory of neural activation. Given the input narrative, we first remove stop words (e.g. the, in, at, that, which, etc.) which provide no semantic information (Silva & Ribeiro 2003). The remaining input words are transformed into word embedding vectors by the Wikipedia2Vec model, pre-trained on the 3 billion word 2018 Wikipeida corpus. These vectors are then input to the reservoir, a recurrent network with fixed recurrent connections. The fixed connections in the recurrent network allow the full dynamics of the system to be exploited, which in some other networks with modifiable recurrent connections is not the case due to temporal cut-off of the recurrence required for implementing learning on the recurrent connections (Elman 1990, Elman 1991, Pearlmutter 1995). This method of fixed connections in the reservoir was first employed in order to model primate prefrontal cortex neurons during behavioral sequence learning tasks (Dominey 1995a), and was subsequently developed with spiking neurons in the liquid state machine (Maass et al 2002), and in the context of non-linear dynamics signal processing with the echo state network (Jaeger 2001, Jaeger & Haas 2004), all corresponding to the class of reservoir computing (Lukosevicius & Jaeger 2009).

In reservoir computing, the principle is to create a random dynamic recurrent neural network, and then stimulate the reservoir with input, and harvest the rich high dimensional states. Typically this harvesting consists in training the output weights from reservoir units to output units, and then running the system on new inputs and collecting the resulting outputs from the trained system. In the current research we focus our analysis directly on the rich high dimensional states in the reservoir itself. That is, we do not train the reservoir to perform any transformation on the inputs. Instead, we analyze the activity of the reservoir neurons themselves. The basic discrete-time, tanh-unit echo state network with *N* reservoir units and *K* inputs is characterized by the state update equation:

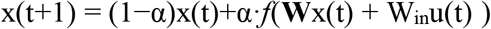

where **x**(*n*) is the *N*-dimensional reservoir state, *ƒ* is the tanh function, **W** is the *N*×*N* reservoir weight matrix, **W***in* is the *N*×*K* input weight matrix, **u**(*n*) is the *K* dimensional input signal, α is the leaking rate. The matrix elements of W and W_in_ are drawn from a random distribution.

The reservoir was instantiated using easyesn, a python library for recurrent neural networks using echo state networks (https://pypi.org/project/easyesn/) (Thiede & Zimmermann 2017). We used a reservoirs of 100 for the simple segmentation experiments, and 1000 neurons for the other simulations, with input and output dimensions of 100. The leak rate was 0.05. This was established based on our empirical observations and the high volatility of the input. Similar results were obtained with leak rate of 0.1. To simulate narrative processing, words were presented in their sequential narrative order to the reservoir. Stop words (e.g. the, a, it) were removed, as they provide no semantic information (Silva & Ribeiro 2003). Similar results were obtained in the presence of stop words. Words were coded as 100 dimensional vectors from the Wikipedia2vec language model. Figure 11 illustrates the forms of the input, and reservoir activation, and their respective cross-correlations.

**Figure 11.**
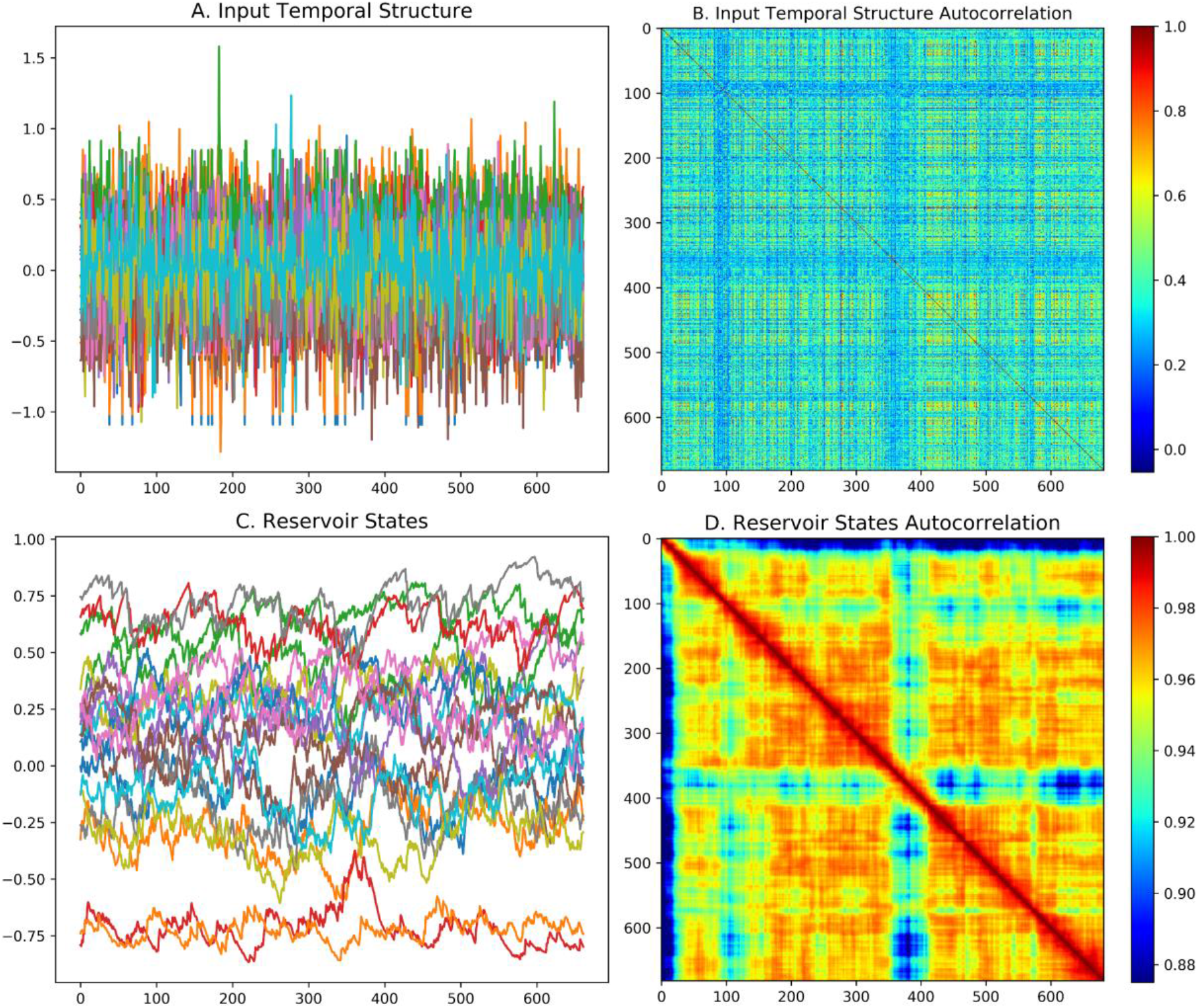
Reservoir fundamentals. A. Temporal structure of input sequence of word embeddings for 100 input dimensions. B. Time-point time-point pattern correlation for input sequence. C. Activity of a subset of reservoir units. Note the relative smoothing with respect to the input in panel A. D. Time-point time-point pattern correlation of reservoir activity during the processing of the narrative. Note the display of coherent structure along and around the diagonal, when compared with panel B. This indicates the integrative function of the reservoir.

In Figure 11, in the upper left panel we see the high frequency of change in the input signal. This is the word by word succession of 100 dimensional word embeddings for the successive words in the narrative. In the upper right we see the auto-correlation, and note that along the diagonal there are not strong correlations. In the lower left we see at the same timescale the activation of 50 reservoir units. Here we can observe that the frequency of change is much lower than in the original input. This is due to the integrative properties of the reservoir. On the lower right we see the autocorrelation of the reservoir states over the time course of the narrative. Here we see along the diagonal more evidence of structure and the integration over time in local patches.

The reservoir has inherent temporal dynamics. We can visualize these dynamics by exposing the reservoir to a zero input, then a constant input, and then return to zero, and then observing the responses to these transitions. Such behavior is illustrated in Figure 12. A zero input is provided from 0 to 500 time steps, then a constant input from 500 to 900, and finally a zero input from 900 to 1500. Figure 12 displays the response of 10 sample neurons. This illustrates the inherent temporal dynamics of the reservoir. In order to more carefully characterize the temporal porperties of the reservoir, we measured the time contants for neural responses to these transitions, and then plotted the ordered time constants. In Figure13 we display the ordered time constants for neurons in response to the transition from zero input to a fixed non-zero signal, and then from signal to zero. These can be compared to the time constants for construction and forgetting.

**Figure 12.**
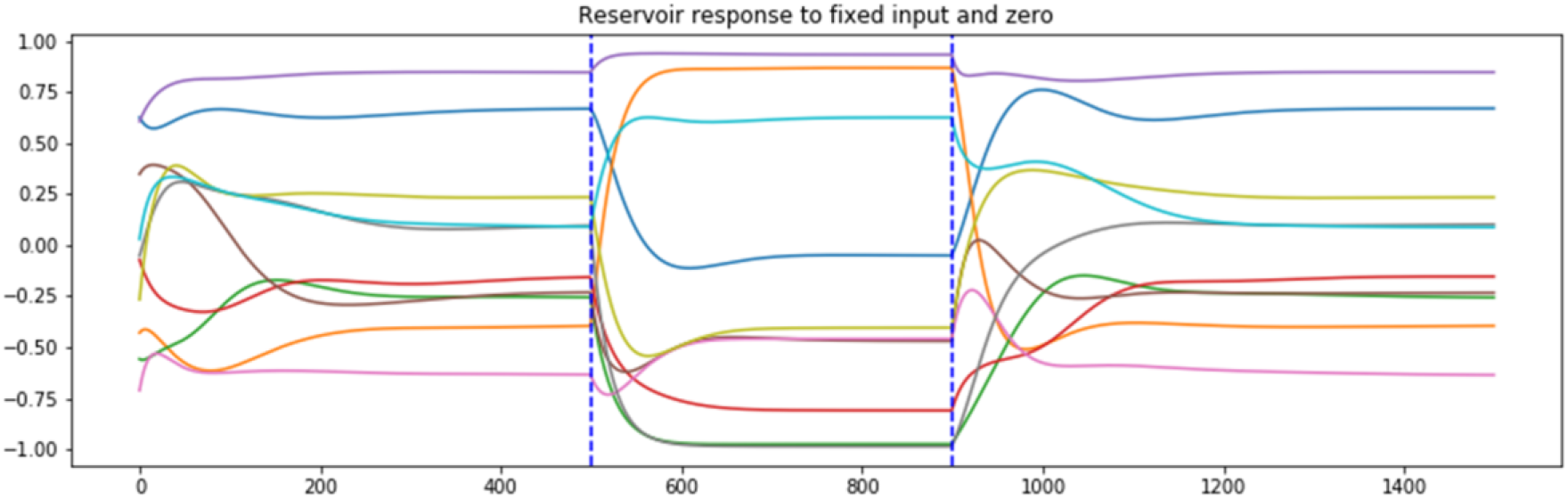
Reservoir dynamics in response to a continuous input of zero, then a fixed non-zero input, and final return to zero input. Note the diversity of temporal responses. This indicates the inherent property of distributed time constants generated by recurrent connections within the reservoir.

**Figure 13.**
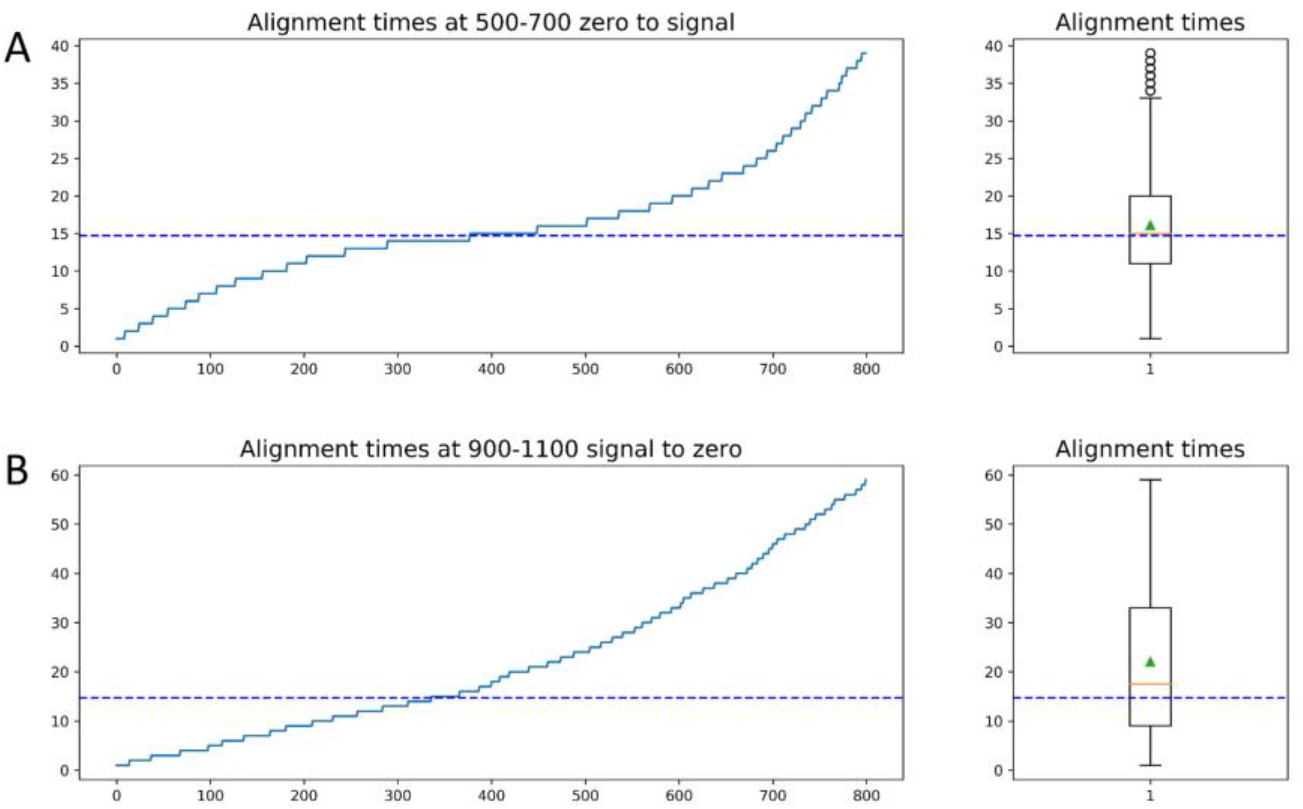
Reservoir units sorted by time to stabilize after the transitions from zero to non-zero input, and from non-zero to zero input. Comparable to Figure 7. Dotted line marks the mean from Figure 7 for comparison.

### Brainiak HMM Model

The HMM model is described in detail in Baldassano et al (2017) and is available as part of the Brainiak python libarary, along with example jupyter notebook that corresponds to the 2017 Baldassano et al. paper^3^. Given a set of (unlabeled) time courses of (simulated or real) neural activity, the goal of the event segmentation model is to temporally divide the data into ‘‘events’’ with stable activity patterns, punctuated by ‘‘event boundaries’’ at which activity patterns rapidly transition to a new stable pattern. The number and locations of these event boundaries can then be compared between human neural activity and simulated Narrative Integration Model activity, and to ground truth values.

The model is run on a given fMRI or reservoir trajectory, and requires specification of the expected number of segments, K. The segmentation returns a probability value. The trained model can then be run on a test trajectory, and a log likelihood score is returned. This mode can be used to evaluate the trained model on untrained test data.

### Model code and data

This research is realized in the open code spirit, and indeed benefitted from open code and data for the fMRI experiments (Chien & Honey 2020, Nastase et al 2020) and the HMM segmentation model (Baldassano et al 2017). The Narrative Integration Model code in python, and all required data is available on Github^4^.

## Acknowledgements

This research was funded by the French Région Bourgogne Franche Comté, Grant ANER RobotSElf.

## Footnotes

1 See Brainiak Tutorial: https://brainiak.org/tutorials/12-hmm/

2 http://datasets.datalad.org/?dir=/labs/hasson/narratives/stimuli

3 https://github.com/brainiak/brainiak/tree/master/examples/eventseg

4 https://github.com/pfdominey/Narrative-Integration-Reservoir/

